# Loss of αBa-crystallin, but not αA-crystallin, increases age-related cataract in the zebrafish lens

**DOI:** 10.1101/2024.01.03.574085

**Authors:** Mason Posner, Taylor Garver, Taylor Kaye, Stuart Brdicka, Madison Suttle, Bryce Patterson, Dylan R. Farnsworth

## Abstract

The vertebrate eye lens is an unusual organ in that most of its cells lack nuclei and the ability to replace aging protein. The small heat shock protein α-crystallins evolved to become key components of this lens, possibly because of their ability to prevent aggregation of aging protein that would otherwise lead to lens opacity. Most vertebrates express two α-crystallins, αA- and αB-crystallin, and mutations in each are linked to human cataract. In a mouse knockout model only the loss of αA-crystallin led to early-stage lens cataract. We have used the zebrafish as a model system to investigate the role of α-crystallins during lens development. Interestingly, while zebrafish express one lens-specific αA-crystallin gene (*cryaa*), they express two αB-crystallin genes, with one evolving lens specificity (*cryaba*) and the other retaining the broad expression of its mammalian ortholog (*cryabb*). In this study we used individual mutant zebrafish lines for all three α-crystallin genes to determine the impact of their loss on age-related cataract. Surprisingly, unlike mouse knockout models, we found that the loss of the αBa-crystallin gene *cryaba* led to an increase in lens opacity compared to *cryaa* null fish at 24 months of age. Loss of αA-crystallin did not increase the prevalence of cataract. We also used single cell RNA-Seq and RT-qPCR data to show a shift in the lens expression of zebrafish α-crystallins between 5 and 10 days post fertilization (dpf), with 5 and 6 dpf lenses expressing *cryaa* almost exclusively, and expression of *cryaba* and *cryabb* becoming more prominent after 10 dpf. These data show that *cryaa* is the primary α-crystallin during early lens development, while the protective role for *cryaba* becomes more important during lens aging. This study is the first to quantify cataract prevalence in wild-type zebrafish, showing that lens opacities develop in approximately 25% of fish by 18 months of age. None of the three α-crystallin mutants showed a compensatory increase in the expression of the remaining two crystallins, or in the abundant βB1-crystallin. Overall, these findings indicate an ontogenetic shift in the functional importance of individual α-crystallins during zebrafish lens development. Our finding that the lens-specific zebrafish αBa-crystallin plays the leading role in preventing age-related cataract adds a new twist to our understanding of vertebrate lens evolution.

## 1. Introduction

Vision is enabled by a properly shaped and transparent lens – a deeply conserved structure found within the camera-eye of groups otherwise highly divergent such as cephalopods and vertebrates. The vertebrate eye lens is primarily composed of proteins from three families: the α, β and γ-crystallins, which contribute to the transparency and refractive power required to focus light onto the retina (Bloemendal & Cate, 1959; Harding & Dilley, 1976; Wistow & Piatigorsky, 1988; Bloemendal & Jong, 1991; Andley, 2007). Surprisingly, α-crystallin gene sequences were found to be similar to the small heat shock protein genes in *Drosophila*, suggesting that these abundant lens materials were evolutionarily coopted from stress-induced protective proteins (Ingolia & Craig, 1982). Vertebrate lenses have two distinct cell-types: fiber and epithelial cells. In fiber cells, the mammalian α-crystallin has been shown to function as a molecular chaperone, preventing the aggregation of denatured protein that could otherwise lead to lens cataract (Horwitz, 1992). This chaperone function is critical for lens homeostasis as lens fiber cells lose their nuclei during development and are not able to replace aging protein (Kuwabara & Imaizumi, 1974; Piatigorsky, 1981; Bassnett, 2002).

Mammals and birds produce two α-crystallin proteins due to a gene duplication event early in vertebrate evolution (Wistow & Piatigorsky, 1988). One of these, αA-crystallin, is primarily expressed in the lens, but its paralog αB-crystallin is widely expressed in lens, nervous and muscular tissue (Bhat & Nagineni, 1989; Dubin, Wawrousek & Piatigorsky, 1989). Mutations in the *cryaa* gene for αA-crystallin can lead to congenital cataract, while those in the *cryab* gene for αB-crystallin can produce both cataract and cardiomyopathies (Litt et al., 1998; Vicart et al., 1998). A variety of α-crystallin mutants and genetic modifications have been used to explore the structure/function relationships of these proteins (Shiels & Hejtmancik, 2021). In mouse knockout models, loss of αA-crystallin led to lens cataract while loss of αB-crystallin did not, suggesting at least in this species that αA-crystallin is more protective against the loss of lens transparency (Brady et al., 1997, 2001).

The zebrafish lens proteome shares many similarities with mammals (Posner et al., 2008; Greiling, Houck & Clark, 2009). However, the genome duplication that occurred at the base of teleost evolution led to the presence of two αB-crystallin paralogs (Postlethwait et al., 1998; Smith et al., 2006). Interestingly, one of these paralogs, αBa-crystallin, has evolved lens-preferred expression while the other, αBb-crystallin, maintains the broad expression found in the single-copy mammalian protein (Posner, Kantorow & Horwitz, 1999; Smith et al., 2006).

The protective chaperone activities of the two αB-crystallin paralogs have also diverged when measured *in vitro*, although there are conflicting data on which is the stronger chaperone (Smith et al., 2006; Koteiche et al., 2015).

Zebrafish are an excellent model system for studying the genetic and molecular processes that guide lens development (Vihtelic, 2008). Their external development, and our ability to perform mutant-analysis, make zebrafish embryos and larvae ideally suited to investigating early events in lens cell-type specification. Mutations of each of the three αB-crystallin genes have been used to examine their impact on early lens development. Zebrafish mutants that do not express αA-crystallin show subtle defects, such as roughness and irregular borders between fiber cells, that were observable using differential interference contrast microscopy (Zou et al., 2015; Posner et al., 2022). Results with αB-crystallin mutants were mixed, with one study showing lens developmental defects while another showed no effect (Mishra et al., 2018; Posner et al., 2022). It is not known how the loss of α-crystallins might affect cataract development as fish age. In fact, little is known about the prevalence of lens cataract in lab-raised zebrafish. Most work on fish lens cataract has focused on aquacultured species, such as salmon and lumpfish (Richardson et al., 1985; Jonassen et al., 2017). Considering the presence of two α-crystallins with lens-preferred expression in zebrafish, it is unclear which of these proteins performs the protective role played by αA-crystallin in the mammalian lens.

The goal of this study was to investigate the role of α-crystallin during lens aging. Zebrafish raised in the lab can live for 3 to 4 years. In this study we examined fish at 6-month intervals through 24 months of age. We hypothesized that the presence of two lens-preferred αB-crystallins in zebrafish might have led to evolutionary changes in their protective roles against cataract formation. The data generated address fundamental questions about the role played by a group of proteins central to the evolutionary success of the vertebrate lens.

## 2. Materials and Methods

### 2.1 Fish maintenance and rearing, study design and data availability

Maintenance and use of zebrafish in this study were approved by Ashland University’s Animal Use and Care Committee (approval #MP 2019-1). Methods of anesthesia and euthanasia followed the *Guidelines for Use of Zebrafish in the NIH Intramural Research Program* and were in accordance with the AVMA guidelines. ZDR strain zebrafish were maintained on a recirculating aquarium system at approximately 28°C with a 14:10 light and dark cycle with daily water changes. Adults were fed twice each day with a combination of dry flake food and live *Artemia* cultures. Adults were bred in false bottom tanks to collect fertilized eggs, which were incubated in petri dishes with system water at 28°C. The resulting larvae that were to be raised to adulthood were fed a liquid slurry of ground zooplankton and green algae until large enough to eat flake food and *Artemia*.

Study design and reporting follows the recommendations in the ARRIVE guidelines (Kilkenny et al., 2010). Any exceptions are noted in the methods below. All quantitative data are included as supplemental tables and all images used to analyze aging lenses are available on Dryad.org.

### 2.2 Single cell RNA-Seq

Fish were maintained by the University of Oregon Zebrafish facility using standard husbandry techniques (Westerfield, 2007). Normal appearing embryos were collected from natural matings, staged and pooled (15 per replicate). Animals used in this study were wild-type (ABC).

#### 2.2.1 Embryo dissociation

As previously reported (Farnsworth, Saunders & Miller, 2020), Collagenase P was prepared to a 100 mg/mL stock solution in HBSS. Chemical dissociation was performed using 0.25% Trypsin, Collagenase P (2 mg/mL), 1 mM EDTA (pH 8.0), and PBS for 15 min at 28C with gently pipetting every 5 min. Dissociation was quenched using 5% calf serum, 1 mM CaCl2, and PBS. Cells were washed and resuspended in chilled (4C), 1% calf serum, 0.8 mM CaCl2, 50 U/mL penicillin, 0.05 mg/ mL streptomycin, and DMEM and passed through a 40 μM cell strainer (Falcon) and diluted into PBS with 0.04% BSA to reduce clumping. A final sample cell concentration of 2000 cells per microliter, as determined on a Biorad TC20 cell counter, was prepared in PBS with 0.04% BSA for cDNA library preparation.

#### 2.2.2. Single-cell cDNA library preparation

Sample preparation was performed by the University of Oregon Genomics and Cell Characterization core facility (https://gc3f.uor <u>egon.edu/</u>). Dissociated cells were run on a 10X Chromium platform using 10x v3 chemistry targeting 10,000 cells. Dissociated samples for each time point for this study (3, 4, 6 and 7 dpf) were submitted in duplicate to determine technical reproducibility. The resulting cDNA libraries were amplified with 15 cycles of PCR and sequenced on an Illumina Hi-seq.

#### 2.2.3. Computational analysis

The resulting sequencing data were analyzed using the 10X Cell-ranger pipeline, version

6.1.2 (Zheng et al., 2017) and the Seurat, version 4.0.6 software package (Satija et al., 2015) for R, version 4.1.2 (Team, 2023a) using standard quality control, normalization, and analysis steps. Briefly, we aligned reads to the zebrafish genome, GRCz11_93, and counted expression of protein coding reads. The resulting matrices were read into Seurat where they were merged with previously obtained data from 1, 2 and 5 dpf larvae (Farnsworth, Saunders & Miller, 2020). The resulting dataset was SC Transformed, and PCA was conducted using 115 PCs. UMAP analysis was performed on the resulting dataset with 115 dimensions and a resolution of 24.0. Based on expression of lens marker genes (*cryaa*, *cryaba*, *cryabb*, *foxe3*, *mipa*), and the presence of cells previously identified as lens fiber and epithelial cells (Farnsworth, Posner & Miller, 2021) we subsetted clusters 394, 411, 197, 347 for further analysis. The cells contained within this lens subset were reclustered using 15 dimensions and a resolution of 0.2. Very few cells from the 7 dpf samples were observed so these samples were excluded from further analysis. The RDS for this subset is available at www.farnsworthlab.com/data, as well as companion code. The raw data will be deposited in the SRA, accession # TBD.

### 2.3 Reverse Transcription Quantitative PCR analysis of lens crystallin expression

Method was designed to meet MIQE guidelines (Bustin et al., 2009). Lenses were removed from zebrafish at four ages to measure ontogenetic changes in mRNA levels of the α-crystallin genes *cryaa*, *cryaba* and *cryabb* using RT-qPCR. Larvae at 10- and 19-days post fertilization (dpf), juveniles at 15 weeks post fertilization (wpf) and adults of various ages were used. Selection of individual fish was not based on any apparent differences or phenotypes, although sick individuals would have been avoided and euthanized. Individuals were anesthetized and both lenses were removed by dissection. Lenses were placed temporarily in phosphate buffered saline (PBS) before storing in RNA Later (Thermo). Stored lenses were kept at either 4°C for up to one week or -20°C for up to one month before purification of total RNA with the Monarch Total RNA Miniprep Kit (NEB). We used approximately 30 lenses at 10 dpf, 10 lenses at 19 dpf, 6 lenses at 5 weeks of age and 4 lenses for adult fish. These sample sizes were determined to produce sufficient RNA for cDNA synthesis. Each age was sampled three times to produce three biological replicates. Lenses were homogenized in DNA/RNA protection buffer with a glass homogenizer and then incubated at 55 °C for 5 minutes with proteinase K following the manufacturer’s protocol. After column purification and on-column DNase treatment, RNA samples were eluted in 50 microliters of nuclease free water and quantified in a Nanodrop One (Thermo).

In a separate set of experiments lenses were removed from adult wild-type zebrafish and adults from our *cryaa* and *cryaba* null mutant lines. Once again, individuals were not selected based on any apparent differences between lines. The wild-type and *cryaa* mutant fish were approximately 18 months of age and the *cryaba* mutant fish were 15 months of age when they were anesthetized and lenses were collected. Total RNA from these lenses was collected using the method described above. Four lenses from two individual fish were used for each biological replicate, with three total biological replicates for each genotype.

We used the ProtoScript II First Strand cDNA Synthesis (NEB) kit to convert purified lens RNA into cDNA using the standard protocol with the d(T)_23_ primer. Fifty nanograms of total RNA was used in each cDNA synthesis reaction in a total volume of 20 microliters. Parallel negative control reactions were run without reverse transcriptase. Synthesized cDNA was stored at -20°C until used. Luna Universal qPCR Master Mix (NEB) was used to amplify 2.0 microliters of each cDNA sample in a 20 microliter total reaction using the standard protocol on an Applied Biosystems StepOne Real-Time PCR system (Thermo). Three technical replicates were used for each cDNA sample, with two replicates for each -RT control and only one reaction for the non-template control. Two endogenous control primers were used (*rpl13a* and *eef1a1l1,* aka *ef1a*). Both genes were shown to have similar expression across zebrafish tissues at different stages (Lang, Wang & Zhang, 2016). All primer sequences used in these experiments are shown in **Supplemental Table 1**. All primers were used at a final concentration of 250 nm in reactions with the following parameters: 95°C hold 1 min; 40 cycles of 95°C for 15 sec and 60°C for 45 sec; fast ramp setting. A melt curve analysis was used to confirm that single products were produced and at least one product from all primers was sequenced to confirm its identity. The Cq value for each reaction was calculated by the OneStep software using default settings. We made the *a priori* decision to exclude any technical replicate that differed from the other two by more than 0.5 Cq. Delta Cq values were calculated in Excel using the average of both endogenous controls and plotted using ggplot2 with R in RStudio (Team, 2023a,b). Statistically significant differences between samples (all run in biological triplicate) were calculated in R using the *anova* and *Tukey HSD* functions.

### 2.4 Lens imaging and analysis

Lenses from adult wildtype fish and each of our three α-crystallin mutant lines were removed at 6, 12, 18 and 24 months to assess any abnormalities. One fish from each tank was genotyped to confirm their identity and fish were not selected based on any apparent differences between individuals. We collected lenses from at least 10 fish of each age, except for 24 months, when at least 15 fish were used. The standard length of each fish was measured by ruler and any opacity of the lens or cornea from the intact eye was noted. Lenses were dissected from anesthetized individuals, placed in PBS and imaged under a dissecting microscope at 40X total magnification over a stage micrometer. Lenses were imaged within 15 minutes of removal and placement in PBS as we often saw a visible optical separation between the lens nucleus and cortex after prolonged incubation in PBS. Observation of lens images was used to identify possible phenotypes. All lens images were then blind scored by removing age and genotype information before scoring. Two individuals blind scored all lenses, with one set of scores presented in Figure 5 of this paper and the other as supplemental data. Statistically significant differences in defect proportions between genotypes at each age was calculated using the “N-1” Chi-squared test (https://www.medcalc.org/calc/comparison_of_proportions.php). The diameter of excised lenses was measured using ImageJ, with the micrometer in each image used for calibration. Analysis of lens size was done in R within RStudio using the *anova* and *Tukey HSD* functions.

## 3. Results

### 3.1 Ontogenetic changes in α-crystallin gene expression

We previously showed that the expression of the αA-crystallin gene *cryaa* is detectable in zebrafish lens fiber cells by 2 days post fertilization (dpf), while mRNA from the two αB-crystallin genes (*cryaba* and *cryabb*) are almost undetectable through 5 dpf (Farnsworth, Posner & Miller, 2021). To more precisely determine the temporal dynamics of gene-expression during lens development, we analyzed single-cell transcriptomic changes in developing lens at 24-hour intervals between 1-6 days post fertilization (dpf). We used the expression of *mipa* to identify fiber cells and *foxe3* expression to identify epithelial cells (**Fig 1A**). Interestingly, there was a noticeable separation between 1 dpf epithelial cells and epithelial cells from later stages, as also found in our previous dataset (Farnsworth, Posner & Miller, 2021). There also appears to be continuous, age-related changes in transcriptomic profile of epithelial cells (**Fig 1A, arrow**). In contrast, fiber cells have a more stable transcriptomic profile after 2 dpf.

**Figure 1.**
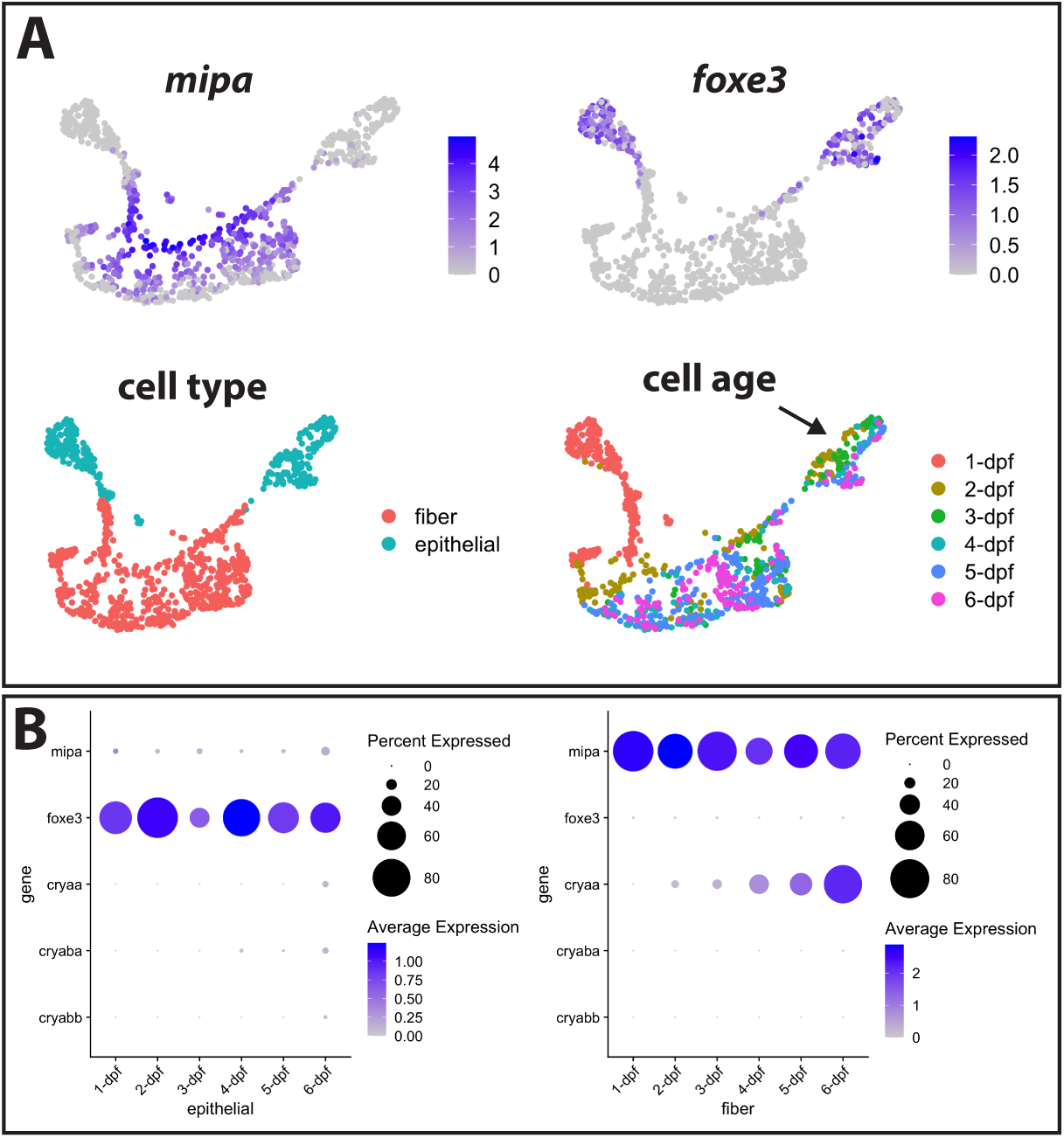
Single cell RNA-Seq analysis of gene expression in zebrafish lens cells through 6 dpf. The expression of genes with known localization to lens fiber cells (*mipa*) and lens epithelial cells (*foxe3*) was used to identify the clusters containing these two cell types (A). Intensity of purple coloration indicates relative expression in each cell. Cells were also colored by age, with an arrow indicating epithelial cells showing a gradient in gene expression between 2 and 6 dpf. The expression of cell type marker genes and the three α-crystallin genes is shown in each cell type from 1 to 6 dpf (B).

Next, we asked if temporal expression for the three α-crystallin genes was a component of age-related changes in early lens development (**Fig 1B**). We previously observed the expression of *cryaa* in fiber cells at 2 dpf. Our analysis in this study revealed increasing *cryaa* expression in an expanding population of fiber cells from 2-6 dpf. In contrast, very little *cryaa* mRNA was detected in epithelial cells until 6 dpf. We once again found very low levels of *cryaba* and *cryabb* expression in both fiber and epithelial lens cells. mRNA levels for *cryaba* did increase slightly in epithelial cells at 4 dpf, and *cryabb* mRNA increases slightly at 6 dpf, but at much lower levels than *cryaa* (**Fig 1B**).

We used a publicly available scRNA-Seq dataset to plot the expression levels of each α-crystallin gene against expression of the lens fiber cell marker *mipa* and epithelial cell marker *foxe3* at 5 and 10 dpf (**Fig 2**) (https://zebrahub.ds.czbiohub.org/; (Lange et al., 2023)). This analysis showed co-expression of *cryaa* with *mipa* at 5 dpf, as expected since *cryaa* also showed fiber cell preferred expression in our scRNA-seq data (**Fig 1**). At this same stage we found no cells co-expressing *mipa* and *cryaba* and only 5 cells co-expressing *mipa* and *cryabb*. At 5 dpf there was very little or no co-expression of any α-crystallin gene with the epithelial cell marker *foxe3*. These expression patterns changed dramatically at 10 dpf. Notably, *cryaa* expression spread to *foxe3* expressing epithelial cells, an increase that also appeared in our new 6 dpf scRNA-seq data (**Fig 1B**). Both αB-crystallin genes also become more broadly expressed in lens cells at 10 dpf, with greater abundance in *mipa* expressing fiber cells (**Fig 2**).

**Figure 2.**
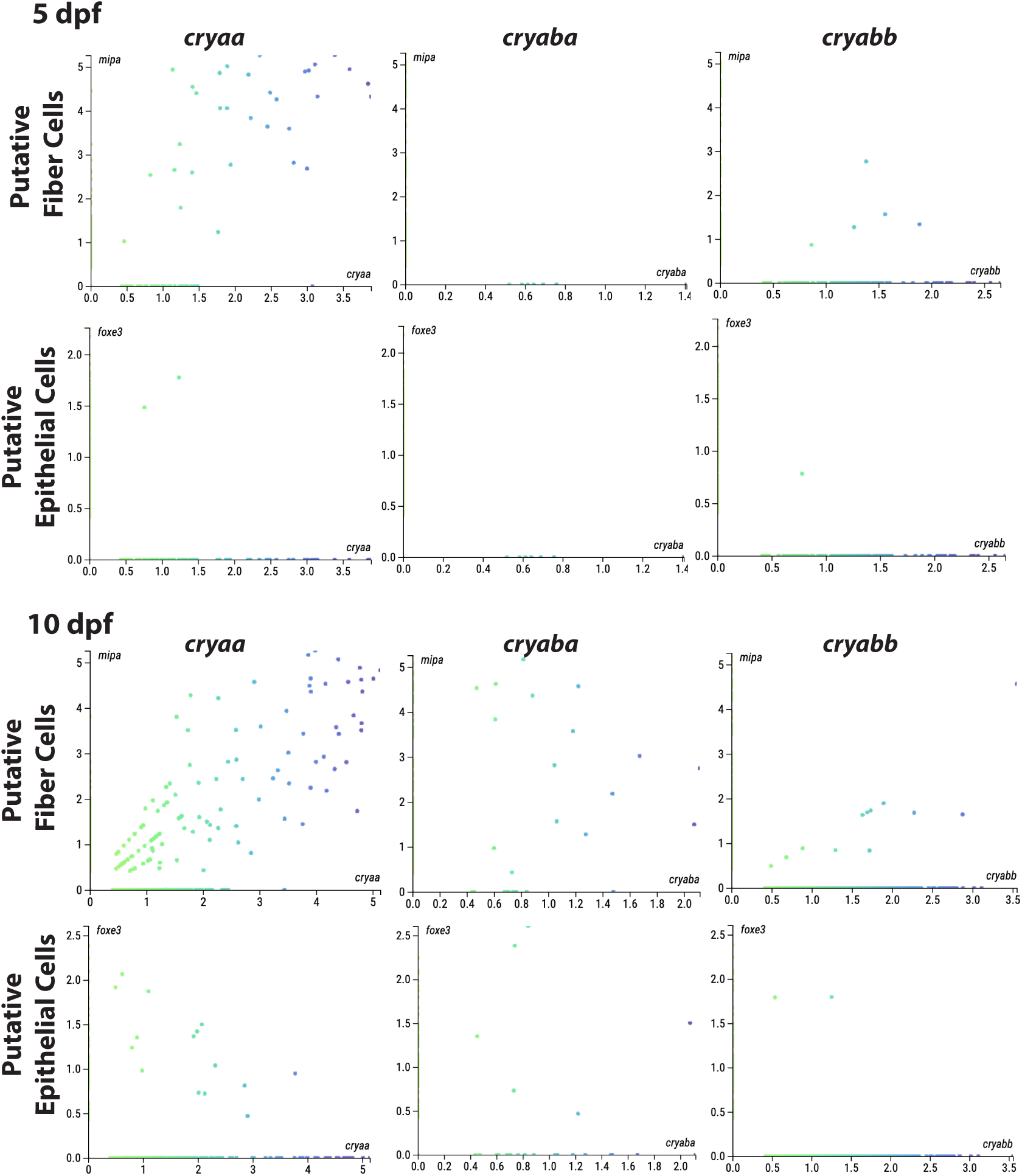
Co-expression of α-crystallin genes with markers for lens fiber and epithelial cells at 5 and 10 days post fertilization based on publicly available single cell RNA-Seq data. The number of cells co-expressing each marker and an α-crystallin gene was used to reflect the levels of expression of that crystallin in each lens cell type. At 5 dpf *cryaa* mRNA was most abundant in putative fiber cells (those cells expressing *mipa*). Putative 5 dpf lens cells co-expressing *cryabb* were much less common, and those with *cryaba* were non-existent. The numbers of putative lens cells co-expressing all three α-crystallin genes increased at 10 dpf, with *cryaa* most abundant. All axes indicate relative transcript counts for the indicated genes. Data are from Zebrahub (https://zebrahub.ds.czbiohub.org/).

There appears to be an increase in the number of lens cells expressing *cryaba* and *cryabb* between 5 and 10 dpf, although this number remains lower than the number of cells expressing *cryaa*.

We used RT-qPCR analysis of RNA from lenses excised from zebrafish at 10 dpf and older to examine ontogenetic shifts in α-crystallin expression as fish age from larvae to juveniles and then adulthood (**Fig 3**). These data showed that *cryaa* lens mRNA was already at its highest levels by 10 dpf, dropped at 19 days, and then increased again to adulthood. By contrast, *cryaba* mRNA levels were lower than *cryaa* levels at 10 dpf, but steadily increased during aging. Lens mRNA for *cryabb* was the lowest of the three at 10 dpf, remained low at 19 days, but increased greatly at 5 weeks and into adulthood (**Fig 3**).

**Figure 3.**
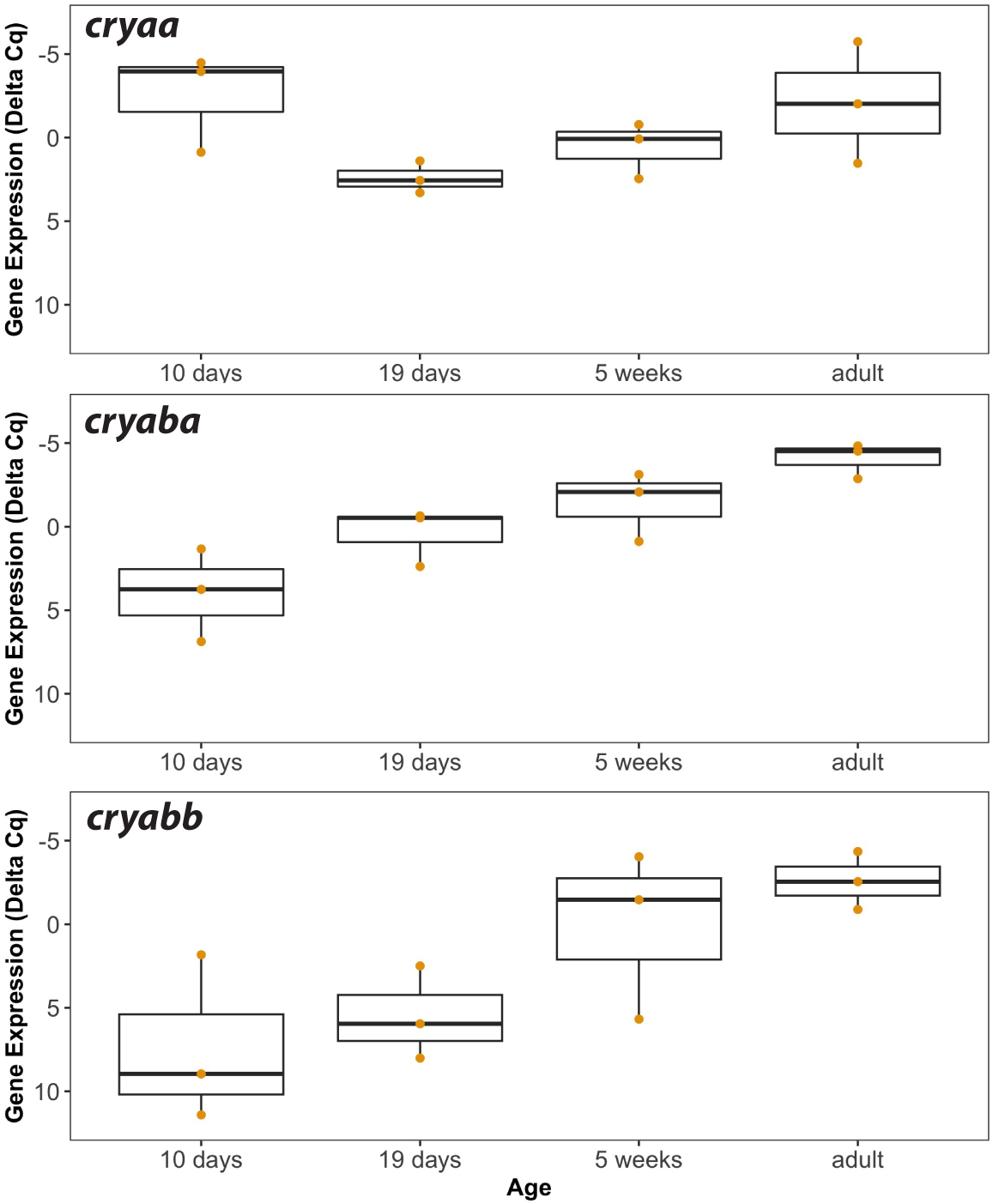
Lens mRNA levels of each α-crystallin gene in larvae (10 and 19 day), juveniles (5 week) and adults. Box and whisker plots are shown for three RT-qPCR biological replicates. The Y-axes are inverted as they show delta Cq values that account for two endogenous control genes (*rpl13a* and *eef1a1l1*), with lower values indicating higher gene expression. Data used for this analysis are included as Supplemental Table 4.

### 3.2 Cataracts and other lens abnormalities in aging zebrafish

We quantified several types of abnormalities in aging zebrafish lenses and compared their abundance between wild-type fish and our three α-crystallin mutants through 24 months post fertilization (mpf). The majority of lenses were clear and produced a sharp image when placed on a stage micrometer (**Fig 4A**). Other lenses showed varied amounts of mild cataract (**Fig 4B**), resulting from both general, uniform cloudiness or opacity in a specific region of the lens. We subjectively further characterized some lenses as having severe cataract (**Fig 4C**). Another distinct lens defect was a dark area found on the anterior pole of the lens (**Fig 4D**), visible through the cornea (**Fig 4E**). Least common were separations between the lens nucleus and cortex and fractures within the lens (**Fig 4F**).

**Figure 4.**
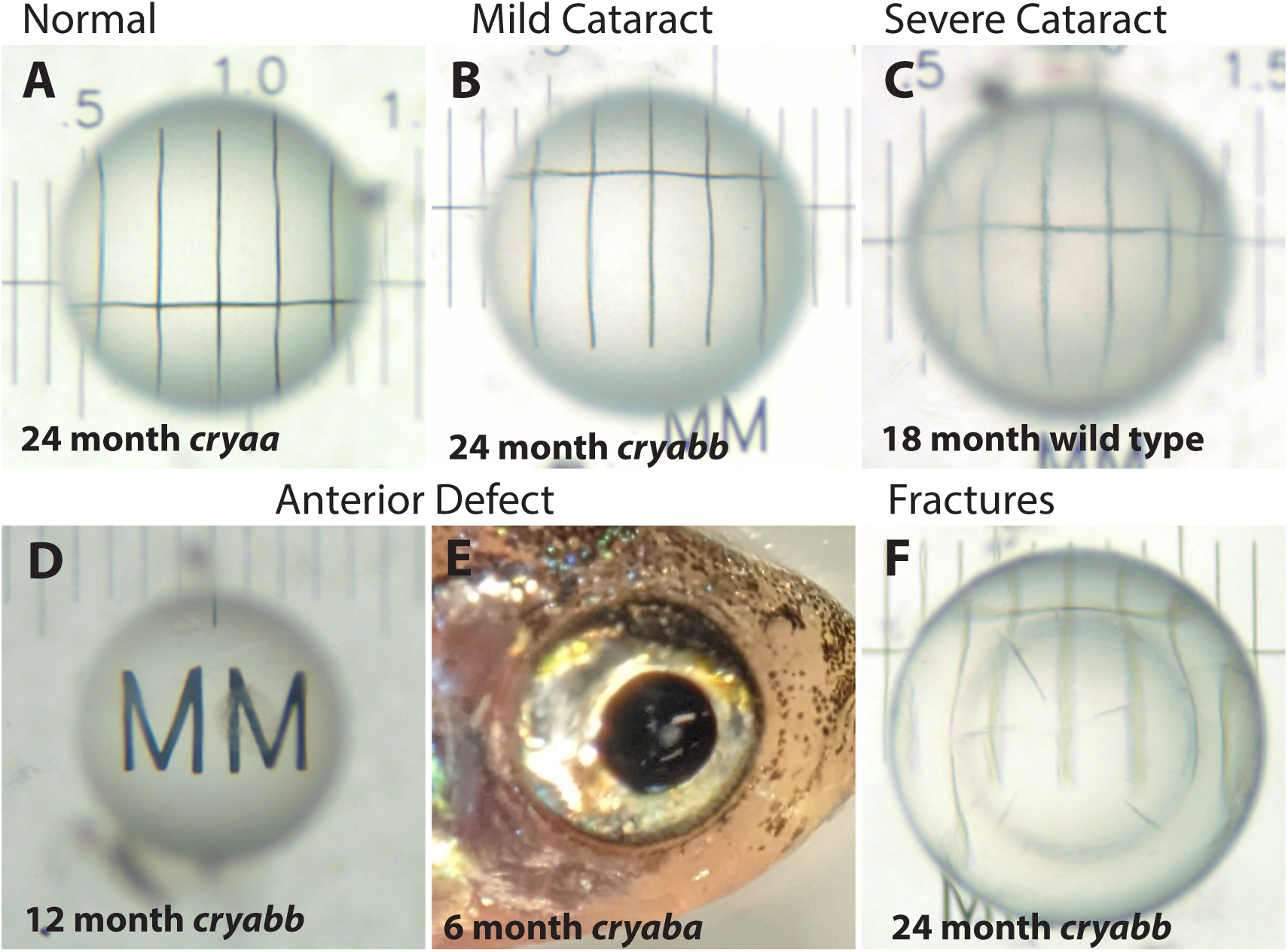
Example images of excised lenses showing typical defects found in wild-type and α-crystallin mutant fish. Lenses were imaged over a stage micrometer to assess any opacity or other defects. Normal lenses were clear and showed sharp micrometer lines across the central portion of the lens (A). Lenses with opacity were blind scored as mild cataract or severe cataract (B and C). Some lenses had a distinct anterior defect (D) that was observable through the cornea before lens removal (E). Some lenses had internal fractures (F). The age and genotype of each example lens is indicated.

**Figure 5.**
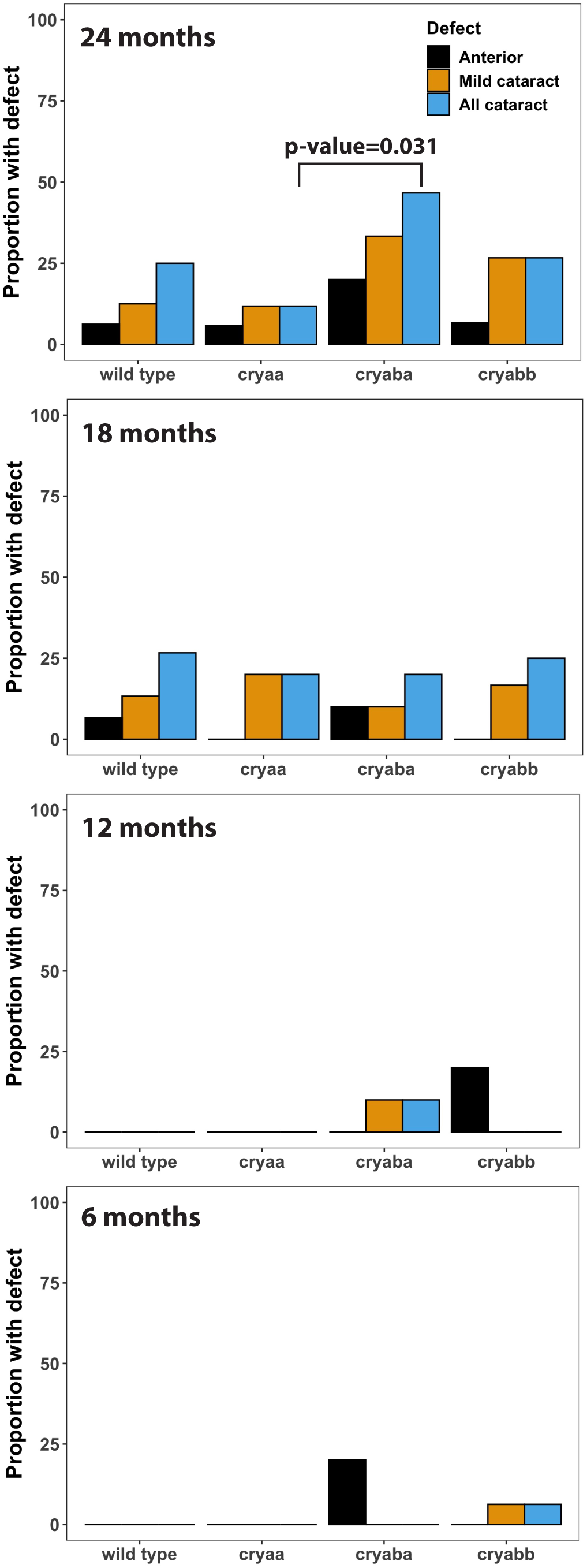
Proportion of wild-type and α-crystallin mutant fish with lens defects from 6 to 24 months of age. Lenses were blind scored with the genetic identities and ages of fish removed to quantify the proportion of lens defects. Any statistically significant differences are indicated with a p-value (“N-1” Chi-squared test). Sample sizes were typically 10 fish for 6, 12 and 18 months and 15 fish for 24 months. Data used for this analysis are included as Supplemental Table 2.

The proportion of wild-type and mutant fishes with each lens defect was quantified by blind-scoring all lens images (**Fig 5**). Lens defects were uncommon at 6 and 12 mpf, but became more common by 18 mpf, when they were fairly uniform across genotypes. We found approximately 25% of wild-type zebrafish had cataract at 18 mpf with no increase at 24 mpf. However, at 24 mpf our *cryaba* mutants showed a higher prevalence of cataract compared to wild type or the other two α-crystallin mutants. This proportion of cataract in *cryaba* mutants was not statistically significantly higher than wild-type fish, but was significantly higher than *cryaa* null mutants (p-value = 0.031). A severe cataract phenotype was also not found in any *cryaa* null mutants, but did occur in wild-type and *cryaba* null mutants. Cataracts occurred both in single lenses in some fish and were bilateral in others. In total, 29 of the 196 examined fish had lens cataract, with 9 of them bilateral. The anterior lens defect occurred in all genotypes, but appeared at younger ages (6 and 12 mpf) only in the *cryaba* and *cryabb* mutants. Male fish made up 64% of our samples, with females 32% and 3.5% not identified. Between genotypes, the proportion of males ranged from 60% in wild-type to 83% in *cryaba* mutants. Of all male fish 17.4% showed cataract while 11% of all female fish showed cataract. This difference is not statistically significantly (p-value 0.2501; “N-1” Chi-squared test). Images of all lenses used in this analysis are publicly available (Dryad.org) and the results of blind scoring are included in Supplemental Table 2.

### 3.3 α-crystallin loss and lens growth

We plotted lens diameter versus fish standard length to visualize whether mutant lenses differed in size from the wild type (**Fig 6A**). It appeared that lenses from *cryaba* mutants tended to be larger than the other genotypes. To quantify this possible difference, we plotted lens diameter as a proportion of standard body length for each genotype at each timepoint. We found no statistically significant differences at 6 and 12 mpf, but at 18 mpf lenses from the *cryaba* mutants were statistically significantly larger than the other genotypes (p-value = 0.020; **Fig 6B**), and both *cryaa* and *cryaba* mutants had larger lenses at 24 mpf (p-value = 0.006 and 0.002, respectively; **Fig 6C**).

**Figure 6.**
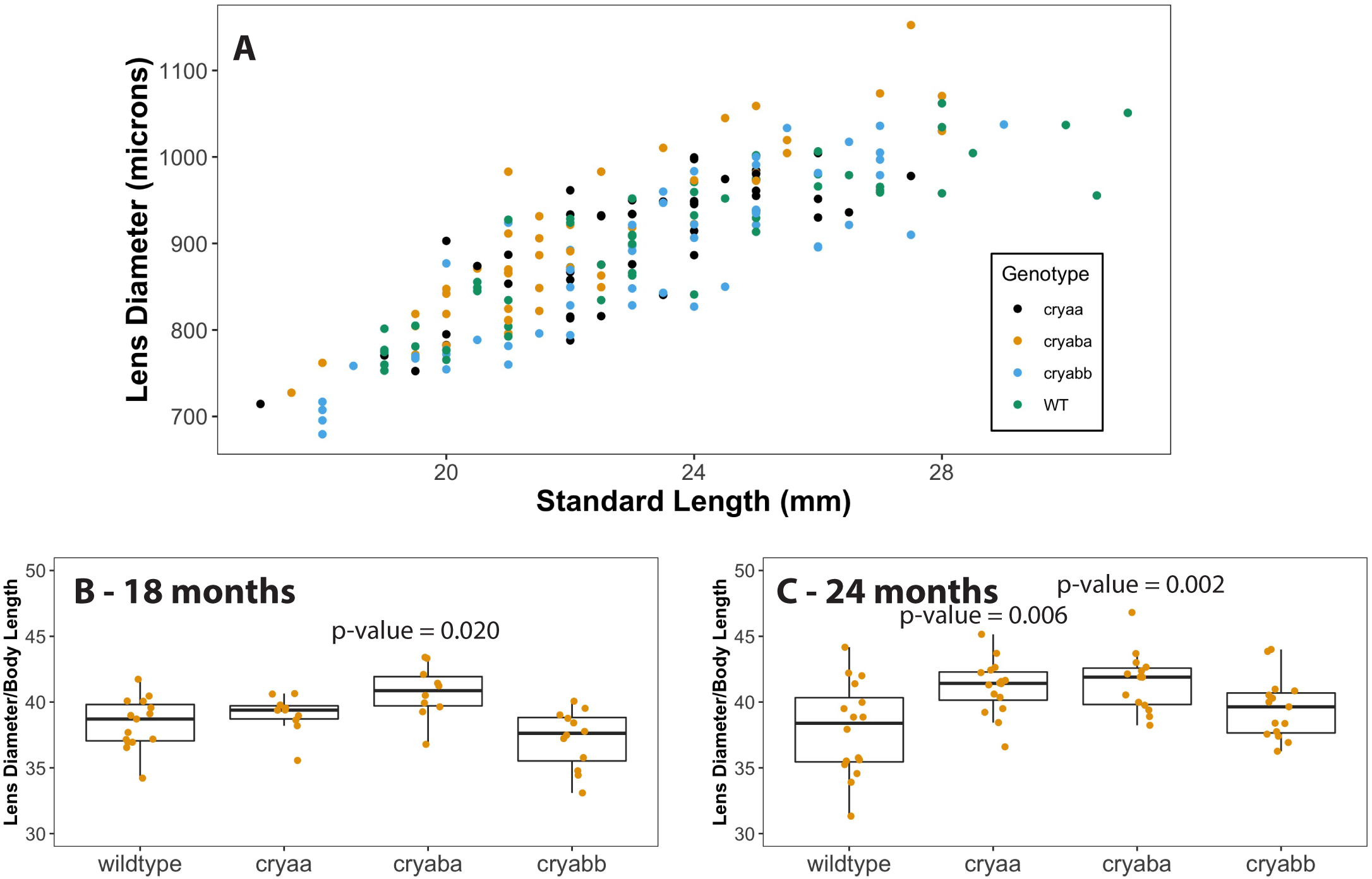
Comparisons of lens diameter to standard length in 18 and 24 month wild-type and α-crystallin mutant fish. Lens diameter (microns) and standard length (millimeters) were measured for each of the four genotypes with the data compiled in one plot (A). Box and whisker plots of that data as ratios for each genotype are shown at both 18 months (B) and 24 months (24). P-values are shown for mutant genotypes that are statistically significantly different than the wild type (ANOVA and tukey post-test). Data used for this analysis are included as Supplemental Table 3.

### 3.4 Lens crystallin expression in adult mutant lenses

We used RT-qPCR to compare transcript levels for the three α-crystallin genes and the most abundant lens crystallin, *crybb1*, between isolated lenses from adult wild-type fish and our *cryaa* and *cryaba* null mutants. We found an expected large reduction in *cryaa* mRNA in our alpha A-crystallin null mutants (**Fig 7A**), as this mutant is missing its proximal promoter and start codon (Posner et al., 2022). While we found a small reduction in *cryaba* transcript in the *cryaba* mutant, this difference was not statistically significant (**Fig 7B**). The *cryaba* mutant still includes its promoter and start codon, and we previously showed that mRNA for this gene was not reduced in whole larvae even though protein was absent (Posner et al., 2022). mRNA levels for *cryabb* and *crybb1* were also similar between wild-type and mutant fish lenses (**Fig 7C and D**). Overall, these RT-qPCR data show that neither the loss of αA-nor αBa-crystallin protein led to changes in the expression of other a-crystallins or the abundantly expressed βB1-crystallin.

**Figure 7.**
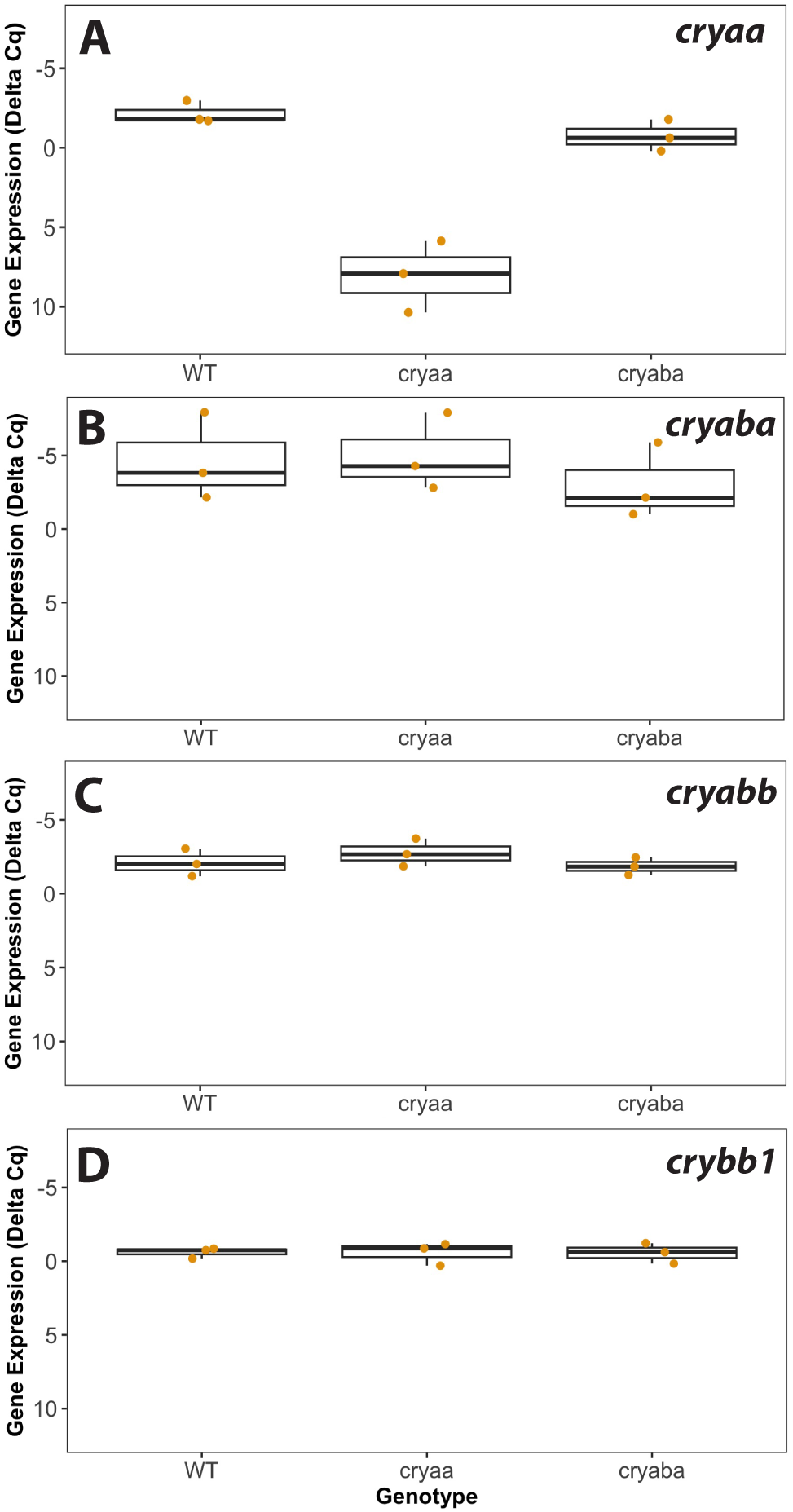
Lens mRNA levels of three α-crystallin genes and the βB1-crystallin gene in wild-type, *cryaa*, and *cryaba* mutant lenses. Box and whisker plots are shown for each crystallin gene, with three RT-qPCR biological replicates for each of the three genotypes. The Y-axes are inverted as they show delta Cq values that account for two endogenous control genes (*rpl13a* and *eef1a1l1)*, with lower values indicating higher gene expression. Data used for this analysis are included as Supplemental Table 5.

## 4. Discussion

The vertebrate lens evolved with two α-crystallin proteins, a lens specific αA-crystallin and ubiquitously expressed αB-crystallin. While these members of the small heat shock protein family both maintain their ability to prevent protein aggregation, it is the lens specific αA-crystallin that is more abundant in the mammalian lens and likely plays a more important role in preventing lens cataract (Brady et al., 1997). That situation appears to have changed in the teleost zebrafish. With the inclusion of a second, lens-specific αBa-crystallin due to the genome duplication at the base of teleost evolution, the zebrafish lens has two α-crystallins with lens preferred expression. We now show that it is αBa-crystallin that plays a more prominent role in preventing lens cataract as the lens ages. These data suggest that the zebrafish, and possibly other teleost fishes, have evolved a second αB-crystallin protein that functions as a protectant against age-induced protein aggregation and cataract. This finding adds an unexpected twist to the interesting history of gene and protein evolution vital for the appearance and function of the vertebrate ocular lens.

While we did not find increased levels of age-related cataracts in fishes lacking αA-crystallin, larvae lacking this protein did show visible lens defects at 3 and 4 dpf (Zou et al., 2015; Posner et al., 2022). Those defects manifested as central roughness, disorganization of central fiber cells, and irregular borders between fiber cells (Posner et al., 2022). The lack of increased cataract in adult *cryaa* null fish suggests that these early developmental defects are either transient, or do not alter transparency as the lens grows. Defects seen in knockout zebrafish for either aquaporin gene *aqp0a* and *aqp0b* were strong at 2 and 3 dpf, but resolved by 4 dpf and did not lead to cataract in adults (Vorontsova et al., 2018). It appears that there can be a disconnect between early developmental phenotypes and the future development of age-related cataracts. The early impact of αA-crystallin loss might be enhanced by the lack of *cryaba* and *cryabb* expression through at least 5 dpf. Our scRNA-Seq data and that from Zebrahub (Lange et al., 2023) both indicate little to no *cryaba* mRNA in lens cells through 5 dpf, with levels increasing at 10 dpf, primarily in fiber cells. If αBa-crystallin has taken over from αA-crystallin as the prominent protectant in the zebrafish lens against protein aggregation, it is possible that any defects produced by αA-crystallin loss are only of significance at early developmental stages.

Previous studies of mouse knockout models for the two mammalian α-crystallins reported changes in lens size and alterations in the transcriptome and proteome. For example, mouse lenses lacking αA-crystallin were smaller in size than wildtype lenses, while loss of αB-crystallin produced no change in lens size (Brady et al., 1997, 2001). We, instead, found a small but statistically significant increase in lens size in our αA- and αBa-crystallin null mutants at 24 months of age. An analysis of the proteome of the mouse αA-/αB-crystallin double knockout did not find a large change in protein expression (Andley et al., 2013). Instead, changes were restricted to increased expression of several histone proteins, β1-catenin, vimentin and βB2-crystallin. Interestingly, not all of these changes at the protein level were reflected in altered mRNA levels. Our finding that mRNA for the three zebrafish α-crystallin genes or the abundantly expressed βB1-crystallin are similar between null mutants and wild-type lenses reflect these earlier findings, and show that transcription of α-crystallin genes does not adjust when αA- or αBa-crystallin protein are missing.

Our RT-qPCR data on adult lenses showed similarly high levels of mRNA for all three α-crystallin genes as well as the abundant βB1-crystallin. Based on past proteomic studies it appears that the amounts of these four proteins differ in the adult zebrafish lens. We previously found that βB1-crystallin made up 7.5% of the adult lens while αBb-crystallin was less than 1% (Posner et al., 2008). Using a rank-based shotgun proteomics approach Greiling et al. (2009) instead found that αA-crystallin was the most abundant protein at 6 months, but again, that αBb-crystallin was least abundant among the four (Greiling, Houck & Clark, 2009). Overall, it appears that these differences in protein levels are not reflected in mRNA concentrations, suggesting that eventual protein amounts are dependent on translational regulation.

We believe that this is the first study to quantify the presence of lens cataracts in wild-type zebrafish during aging. Cataract was not found in 6 and 12 month old fish, but appeared and stayed steady from 18 to 24 months, peaking at approximately 25%. This proportion is far lower than reported for the lumpfish, which has been used to study cataract development in aquaculture (Jonassen et al., 2017; Paradis et al., 2019). However, the presence of some baseline cataract in wild-type zebrafish allows for comparison with mutant strains to determine any possible statistically significant increase in lens opacity. We concluded our data collection at 24 months, earlier than the typical laboratory lifespan of the zebrafish, which can be up to 3.5 years or more. It is very possible that cataract prevalence would increase further beyond 24 months of age.

In summary, this is the first study to quantify the prevalence of cataract in aging zebrafish and determine the impact of α-crystallin loss. Unlike past studies with mouse knockouts, it was the loss of αBa-crystallin, not αA-crystallin, that increased the development of age-related cataracts. Loss of each α-crystallin did not lead to compensatory changes in the transcription of the others. These results are important for future studies using zebrafish as a model for age-related cataract and suggest that in the zebrafish, and perhaps other teleost fishes, the lens specific αBa-crystallin paralog has become the key protein protecting against lens opacity that results from aggregation of aging protein.

## Acknowledgments

This work was supported by the NIH National Eye Institute (R15 EY13535) to MP and by the AU Choose Ohio First program that provided summer research support to KM.

## Author Contributions

MP conceived and designed the study. DRF designed and conducted the scRNA-Seq experiments. MP, TK, SB, MS, TG and BP contributed to the analysis of aging lenses and the RT-qPCR gene expression analysis. MP, TG and DRF wrote the manuscript. All authors reviewed the manuscript.

## Data Availability

Data used for Figures 3, 5, 6 and 7 are included in Supplemental Tables 2-5. Lens images will be made available on Dryad.org. The RDS for the subset of analyzed scRNA-Seq data is available at www.farnsworthlab.com/data, as well as companion code. The raw data will be deposited in the SRA, accession # TBD.

